# Resolution and co-occurrence patterns of *Gardnerella leopoldii*, *Gardnerella swidsinskii*, *Gardnerella piotii* and *Gardnerella vaginalis* within the vaginal microbiome

**DOI:** 10.1101/702647

**Authors:** Janet E. Hill, Arianne Y.K. Albert, the VOGUE Research Group

## Abstract

**Background:** *Gardnerella vaginalis* is a hallmark of vaginal dysbiosis, but is found in the microbiomes of women with and without vaginal symptoms. *G. vaginalis* encompasses diverse taxa differing in attributes that are potentially important for virulence, and there is evidence that ‘clades’ or ‘subgroups’ within the species are differentially associated with clinical outcomes. The *G. vaginalis* species description was recently emended, and three new species within the genus were defined (*leopoldii*, *swidsinskii*, *piotii*). 16S rRNA sequences for the four *Gardnerella* species are all >98.5% identical and no signature sequences differentiate them.

**Results:** We demonstrated that *Gardnerella* species can be resolved using partial chaperonin-60 (cpn60) sequences, with pairwise percent identities of 87.1-97.8% among the type strains. Pairwise co-occurrence patterns of *Gardnerella* spp. in the vaginal microbiomes of 413 reproductive aged Canadian women were investigated, and several significant co-occurrences of species were identified. Abundance of *G. vaginalis*, and *swidsinskii* was associated with vaginal symptoms of abnormal odour and discharge.

**Conclusions:** cpn60 barcode sequencing can provide a rapid assessment of the relative abundance of *Gardnerella* spp. in microbiome samples, providing a powerful method of elucidating associations between these diverse organisms and clinical outcomes. Researchers should consider using cpn60 in place of 16S RNA for better resolution of these important organisms.

## Background

Since its original isolation from human vaginal samples in 1953 [1], the species that eventually became known as *Gardnerella vaginalis* has been strongly associated with vaginal dysbiosis and negative reproductive outcomes [2]. Evaluation of the abundance of *Gardnerella* morphotypes in Gram stained vaginal smears is a key factor in calculation of the Nugent score [3] and thus in the microbiological definition of bacterial vaginosis (BV). Understanding of its role in the vaginal microbiome has been complicated due to phenotypic diversity within the taxon and its perplexing presence, sometimes in high numbers, in the vaginal microbiomes of women without any signs or symptoms of dysbiosis [2].

Over the past decades, several classification schemes have been developed in an attempt to delineate subgroups within *Gardnerella vaginalis*. These have included both “biotyping” schemes based on a set of biochemical test results [4, 5], and molecular methods based on amplification and restriction digestion of 16S rRNA gene sequences [6]. In 2005, four clusters of *G. vaginalis*-like sequences were observed in vaginal microbiome profiles based on sequencing of amplified cpn60 barcode sequences [7]. More recently, four “subgroups” or “clades” of *G. vaginalis* were defined based either on partial sequences of the cpn60 barcode sequence [8–10] or the concatenated sequences of 473 genes common to a set of 17 *G. vaginalis* isolates [11]. When we compared these latter two approaches directly, they were consistent with each other, with cpn60 defined subgroups A-D corresponding to clades 4, 2, 1 and 3, respectively [8]. Based on the results of this comparison, we suggested that pairwise average nucleotide identity (ANI) values between whole genome sequences of representative isolates were consistent with their definition as separate species [12] but that additional phenotypic characteristics that differentiate the subgroups should be identified [8].

In 2019, Vaneecoutte et al. [13] formally proposed the emendment of the species *G. vaginalis*, and defined three new species: *G. piotti*, *G. swidsinskii* and *G. leopoldii*, based on comparison of whole genome sequences, biochemical properties and matrix-assisted laser desorption ionization time-of-flight (MALDI-TOF) mass spectrometry analysis. The species could not be resolved using 16S rRNA gene sequences [13]. In addition to these four species, the authors also defined nine additional “genome species” based on whole genome sequence comparisons. These additional genome species were not named or formally described, presumably due to a lack of sufficient numbers of isolates to make a strong case for their designation.

Given the apparent significance of *Gardnerella* spp. in the vaginal microbiome, resolution of these species in metagenomic samples and association of their presence and abundance with clinical outcomes is critical. We have already demonstrated that amplification and sequencing of the cpn60 barcode can be used to resolve cpn60-defined subgroups A-D in vaginal samples, and demonstrated that their differential abundance can be used to reveal previously unreported community state types [14]. The objectives of the current study were to determine if cpn60 barcode sequences could differentiate the newly defined species and genome species of *Gardnerella*, to investigate their distribution in a collection of 417 previously sequenced vaginal microbiome profiles, and to identify associations of *Gardnerella* spp. with vaginal symptoms. Our results confirm the resolving power of the cpn60 barcode sequence and reveal significant co-occurrences of *Gardnerella* spp. in the vaginal microbiome that have implications for diagnostics for women’s health, and for our understanding of vaginal microbial ecology.

## Methods

### Gardnerella cpn60 sequence analysis

cpn60 universal target sequences from 52 *Gardnerella* species representing the four named species (*G. vaginalis*, *G. piotti*, *G. swidsinskii* and *G. leopoldii*) and nine additional genome species described by Vaneechoutte et al. [13] were retrieved from cpnDB (www.cpndb.ca, [15, 16]) and aligned using CLUSTALw. Inter-species nucleotide sequence similarities were calculated using *dnadist* within PHYLIP [17]. A bootstrapped phylogenetic tree was calculated using *seqboot*, *dnadist* (maximum likelihood option), and *neighbor*. The type strain of *Alloscardovia omnicolens* (DSM 21503) was included as an outgroup. The consensus tree was computed with *consense* and branch lengths applied with *fitch*. The tree vas visualized using FigTree (v1.4.2).

### *Identification of Gardnerella* spp. *in vaginal microbiomes of Canadian women*

cpn60 barcode sequence data, Nugent scores and self-reported symptom data from previously conducted studies by the VOGUE Research Group of the vaginal microbiome composition of non-pregnant, reproductive aged Canadian women recruited from clinics in greater Vancouver, Canada area were used in the current sub-analysis. These included healthy women (n=310), women living with HIV (n=54) and women who had at least four self-identified episodes of vulvovaginitis in the past 12 months (n=53). DNA extraction, cpn60 barcode PCR, library preparation and sequencing of amplicons are described in [14]. Amplification primer sequences were removed using cutadapt, followed by quality trimming with trimmomatic (quality cut-off 30, minimum length 150). Quality trimmed reads were loaded into QIIME2 [18] for sequence variant calling and read frequency calculation with DADA2 [19] and a truncation length of 250. For taxonomic identification, variant sequences were compared to the cpnDB_nr reference database (version 20190305, downloaded from www.cpndb.ca) using watered-BLAST [20]. The reference database includes representative sequences of each of the four named *Gardnerella* species and the nine additional genome species defined by Vaneechoutte et al. [13].

### Co-occurrence and cluster analysis

For the following analyses we removed four samples with total sequence reads <500, leaving 413 samples. We determined pairwise co-occurences using the presence or absence of each *Gardnerella* species or genome species in each sample. Presence was indicated if the raw number of sequence reads in the sample was ≥10, otherwise the species was marked as absent. We calculated a Jaccard index of similarity (*J*) [21] for each pairwise combination and compared these to a probabilistic null model of species co-occurrence that takes into account observed frequencies to determine significance [22, 23](Supplemental File 1). P-values were Benjamini-Hochberg (BH) corrected to a false-discovery rate = 0.05 [24]. Jaccard dissimilarities (1-*J*) were used to cluster the species using complete-linkage hierarchical clustering implemented in the *vegan* package for R [25].

### Comparisons of clinical data and relative abundance

Relative abundances were calculated using the centre-log-ratio transformation as implemented in the *ALDEx2* package [26] and *propr* package [29]. We compared clr abundance among categories of clinical variables using Kruskal-Wallis tests. Significant omnibus tests were followed up with Dunn’s post-hoc tests with BH adjusted p-values. For this analysis, the sample size was reduced to 395 for which we had concurrent Nugent scoring data, and 393 for which we had self-report symptom data.

## Results and Discussion

### Resolution of *Gardnerella* spp. based on cpn60 universal target sequences

The four named *Gardnerella* spp. were resolved in the cpn60 phylogenetic tree based on an alignment of the 552 bp “universal target” sequence barcode (Figure 1), with good bootstrap support for nodes separating the species. *G. vaginalis* (genome sp. 1) corresponds to the previously described subgroup C/clade 1. This observation is consistent with previous demonstrations that cpn60 barcode sequences are generally excellent predictors of whole genome sequence relationships among closely related bacteria [27, 28]

**Figure 1.**
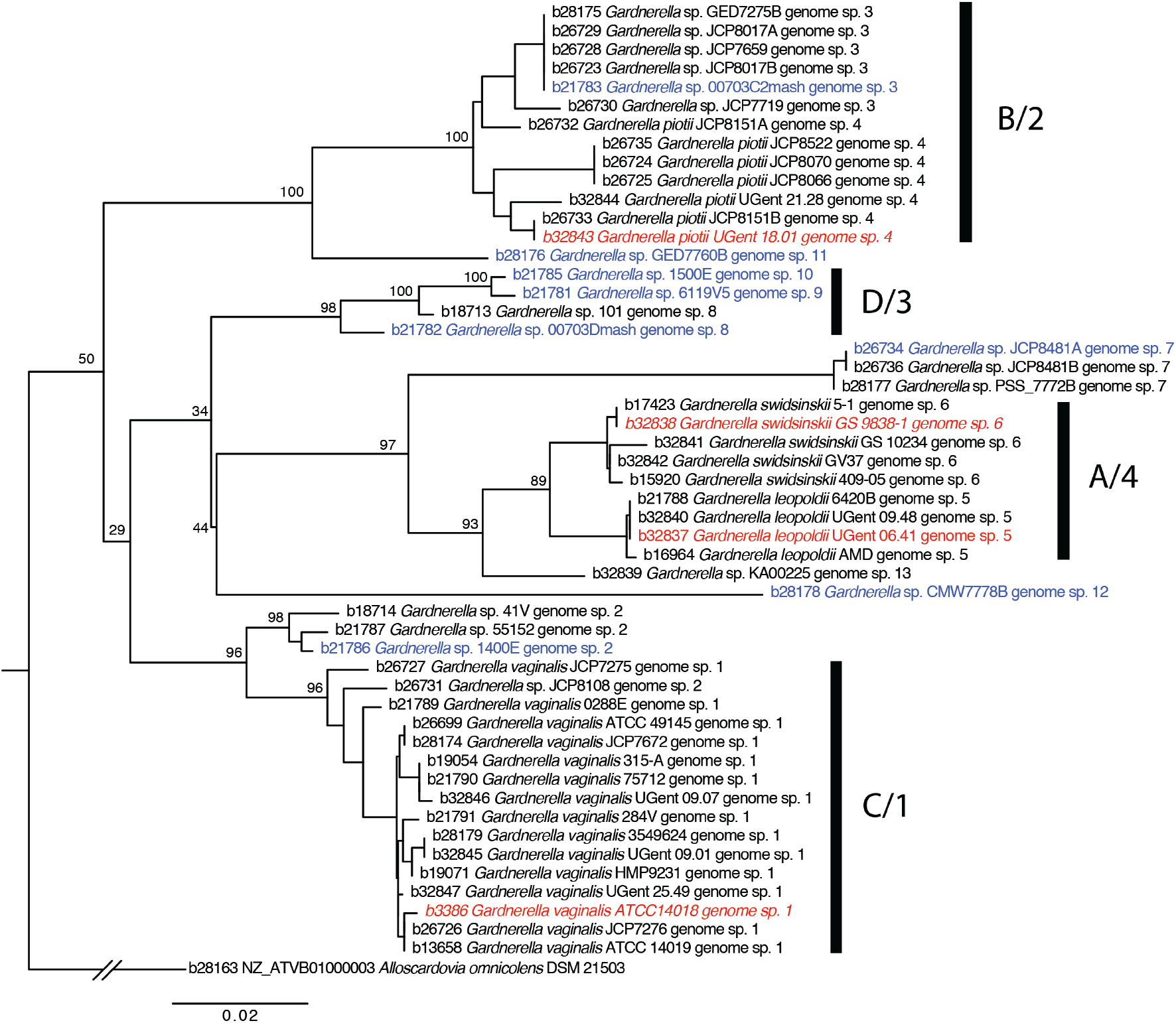
Phylogenetic relationships of *Gardnerella* spp. based on an alignment of the 552 bp *cpn*60 barcode sequence. Type strains are shown in red, and representatives of the other 9 genome species designated by Vaneechoutte et al. [13] are in blue. Bootstrap values are indicated at the major nodes. The tree is rooted with *Alloscardovia omnicolens*. cpn60 subgroups A-D [10], and clades 1-4 [11] are labeled as subgroup/clade based on sequences common to previous studies.

*G. swidsinskii* (genome sp. 6) and *G. leopoldii* (genome sp. 5) share a common node in the tree, but are separated clearly with good bootstrap support. These two species (represented by strains AMD and 5-1) were previously grouped together within cpn60 subgroup A/clade 4 based on the selection of isolates available for analysis at the time. Vaneechoutte et al. noted in their description of the species definitions that *G. swidsinskii* and *G. leopoldii* could be distinguished based on ANI, DNA-DNA hybridization (DDH) and MALDI-TOF profiles [13].

*G. piotii* (genome sp. 4) corresponds to subgroup B/clade 2 and can be distinguished from other species based on ANI, DDH, MALDI-TOF and a positive sialidase test. One *G. piotii* isolate (JCP8151A) clustered with genome sp. 3 in the tree. In the original description of the cpn60 subgroups, what was designated genome sp. 3 by Vaneechoutte et al. was included in subgroup B and it is noteworthy that several isolates examined by Schellenberg et al. have cpn60 sequences identical to strain 00703C2mash (genome sp. 3) were also found to be sialidase positive [8]. The characterization of additional isolates of genome sp. 3 and *G. piotii* will be necessary to determine if these groups should be combined.

Subgroup D/clade 3 was the most diverse in previous descriptions and so it is not surprising to find that isolates previously identified as Subgroup D (strains 101, 1500E, 6119V5 and 00703Dmash) are separated into three genome species: genome spp. 8, 9 and 10. Complete characterization and possible naming of these three genome species, along with genome spp. 2, 7, and 11-13 will require analysis of additional isolates to establish differentiation by whole genome sequence and additional phenotypic characteristics.

Pairwise nucleotide sequence identities for the type strains of *Gardnerella* spp. and representatives of the other nine genome species were calculated from the aligned sequences (Table S1). Identities among the four type strains were from 87.1% (*G. leopoldii* vs. *G. piotii*) to 97.8% (*G. leopoldii* vs. *G. swidsinskii*). When representatives of the other nine genome species were included, percent identities for the 552 bp cpn60 barcode sequence ranged from 84.2% (genome sp. 7 vs. genome sp. 2 or *G. piotii*) to 99.4% (genome sp. 9 vs. genome sp. 10). No isolates had identical cpn60 barcode sequences. Inter-specific cpn60 barcode sequence identities are known to vary widely among bacteria genera so this range was not unexpected [15]. To investigate the resolving power of the 5’ and 3’ ends of the barcode sequence that would be determined using routine next-generation sequencing protocols, we truncated the alignment to examine 250 bp of either end of the barcode sequence. Average pairwise identities were 88.2% (range 83.2 - 99.6) and 91.3% (range 84.4 - 99.2), respectively (Table S1). None of the species were identical. These identities cover the same range as observed for the entire barcode sequence, as is expected given the uniform distribution of sequence variation along its length [9].

### Classification of *Gardnerella* sequence variants

One of the major advantages of use of the cpn60 barcode sequence for taxonomic profiling of microbial communities is the ability to achieve species level classification of sequence reads or assembled operational taxonomic unit (OTU) sequences routinely. It was this resolution that led to the identification of previously undescribed community state types in the human vaginal microbiome, based on the detection of subspecies level sequences within *Gardnerella* [14]. Elucidation of the role of genomically and phenotypically distinct *Gardnerella* lineages in the vaginal microbiome and determining their association with clinical outcomes requires determining their distribution in clinical cohorts. Accomplishing this on a large scale requires culture-independent techniques. While whole-genome shotgun metagenomics might provide resolution of *Gardnerella* spp., this approach requires orders of magnitude more sequencing effort and much more complex bioinformatics than amplicon sequencing. Based on the successful resolution of 13 genome species of *Gardnerella* described above, we next sought to discover if they could be reliably detected and quantified in cpn60 amplicon sequence-based microbiome profiles.

cpn60 barcode sequence data was available from 417 previously characterized vaginal samples from non-pregnant, reproductive aged Canadian women. For the purposes of the current study, exact sequence variants were identified using DADA2 and a truncation length of 250 bp and variants were compared to the cpnDB_nr reference database [15] to identify the nearest database neighbour.

Most (301/413) of the samples for which at least 500 reads were available contained some *Gardnerella*-like sequence variants and variants corresponding to all 13 *Gardnerella* spp. and genome species were detected. The median sequence identity of variants to reference sequences was 98.4%. Sample prevalence and proportional abundance ranged widely among species (Table S2). For example, 68.4% (206/301) of *Gardnerella* positive samples contained *G. vaginalis* and 49% (148/301) contained *G. swidsinskii*, but seven genome species (2, 7–13) were detected in ≤10% of samples. The prevalence and abundance patterns are generally consistent with previous descriptions of vaginal microbiomes based on cpn60 barcode sequencing [14, 29–32] or clade-specific quantitative real-time PCR [33, 34]. There were 60 samples with at least 50% of their read counts accounted for by *Gardnerella* spp., and 30 samples with at least 75% *Gardnerella* (Table S2).

The number of *Gardnerella* spp. detected per sample ranged from one (109/301, 36.2%) to ten (3/301, 1%), although the majority (184/301, 61.1%) contained one or two species (Figure 2). Overall, multiple *Gardnerella* spp. were detected in 63.8% of samples, consistent with a previous report of multiple *Gardnerella* “clades” in 70% of samples from women with BV [33]. The prevalence and proportional abundance of *Gardnerella* spp. in samples with >50% *Gardnerella* (n = 60) are shown in Figure 3. In addition to the four named species, genome sp. 3 was detected frequently and in relatively high proportional abundance, in striking contrast to the rarely detected genome species 2, and 7-13.

**Figure 2.**
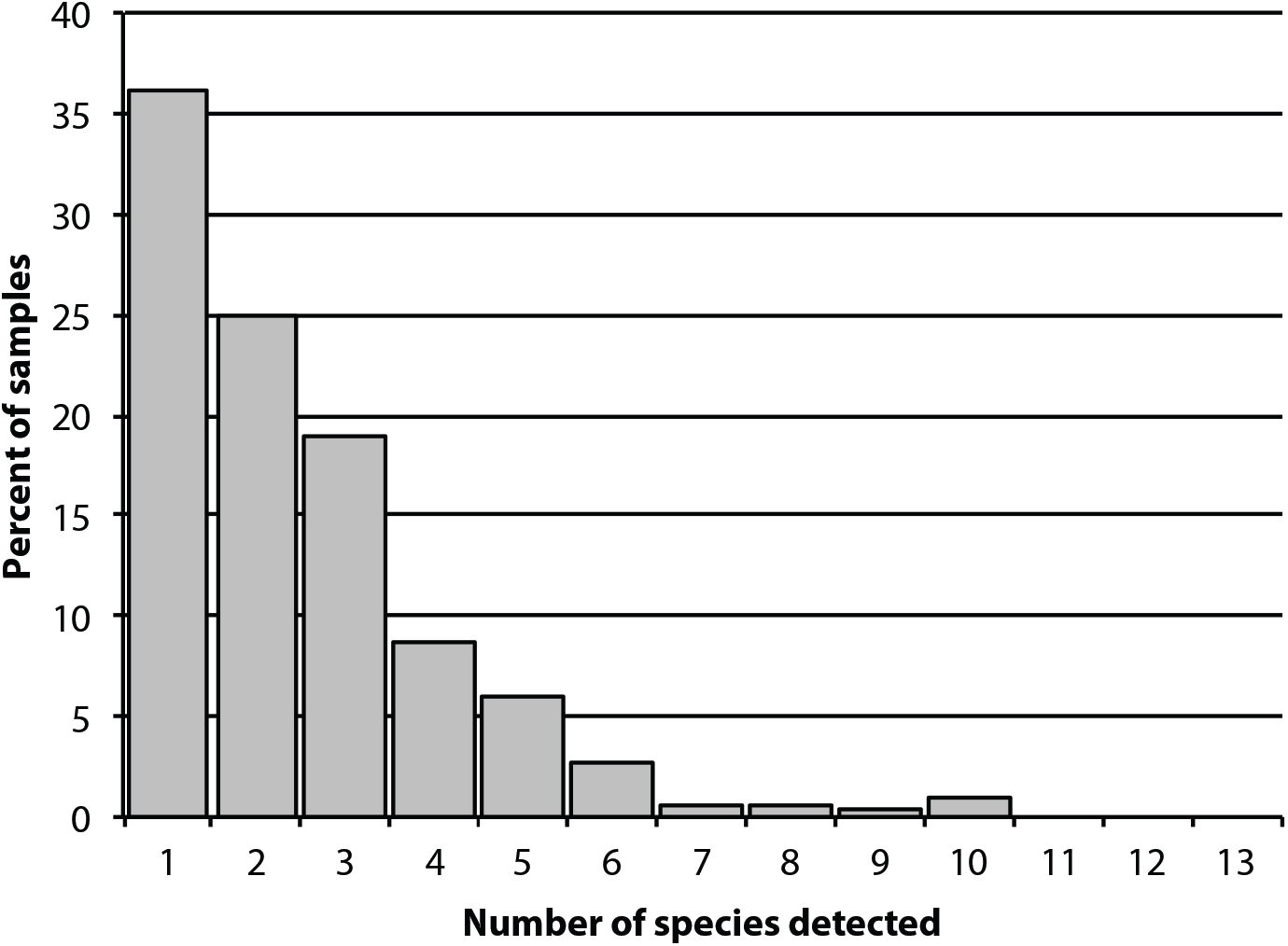
Number of *Gardnerella* spp. detected per sample (n = 301).

**Figure 3.**
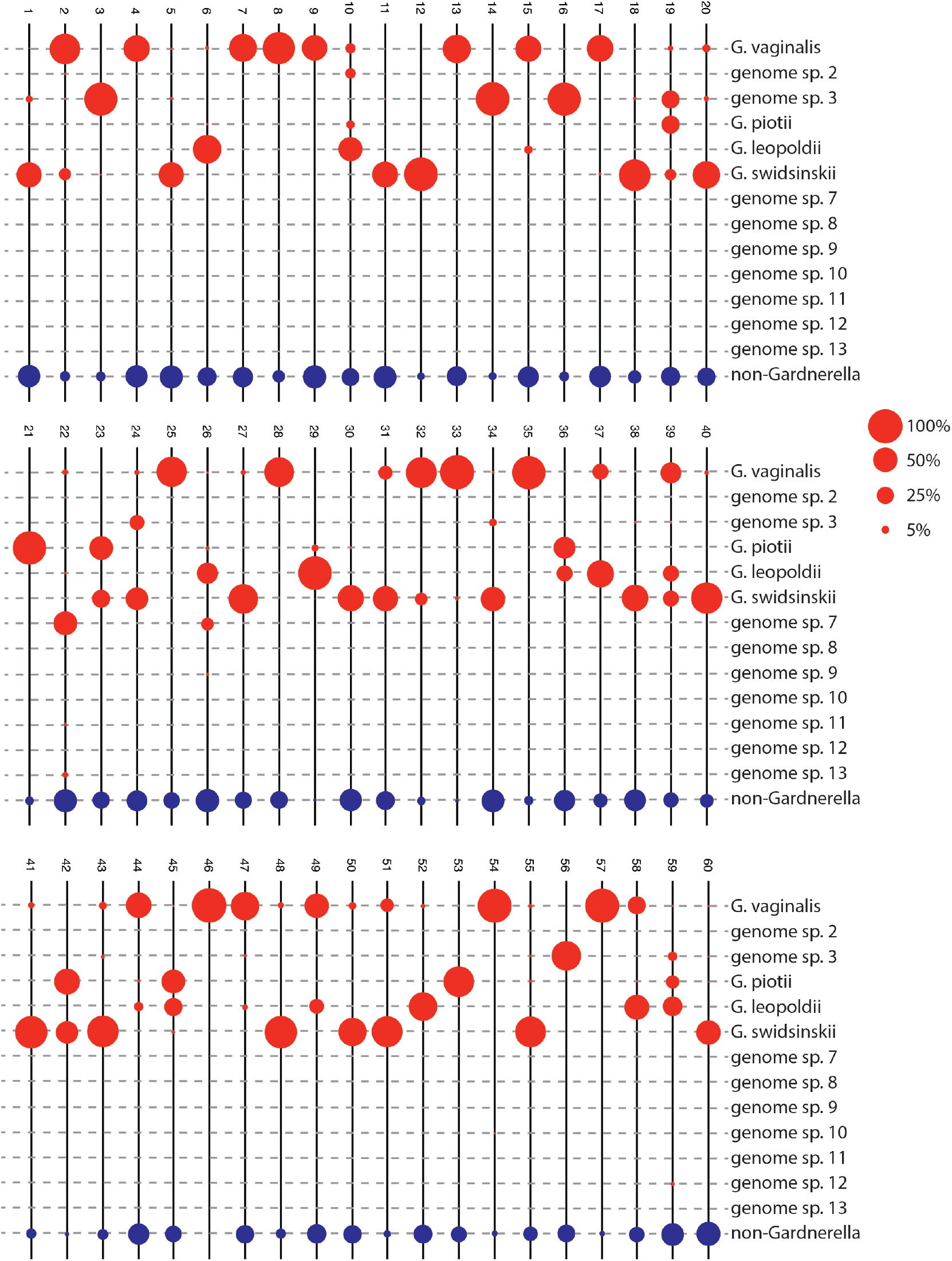
Proportional abundance of 13 *Gardnerella* spp. in vaginal microbiomes of women with ≥50% *Gardnerella* sequence reads (n = 60). Red circles indicate proportional abundance of each species according to scale on the left; blue circles represent the proportion of reads identified as non-*Gardnerella*.

The prevalence and abundance patterns we observed in these samples mirrors the isolate and whole genome sequence collection used to provide evidence for the emendment of *Gardnerella* and the designation of the new species [13]. Based on our experience, there is no obvious bias in PCR amplification of *Gardnerella* lineages, and multiple representatives of all previously defined cpn60 subgroups were readily amplified using cpn60 “universal” PCR primers [8]. Furthermore, we have shown a strong correlation between *Gardnerella* cpn60 sequence read counts in amplicon-based microbiome profiles and abundances determined by *Gardnerella*-specific quantitative real-time PCR [29]. Thus, it seems that some *Gardnerella* species are actually less prevalent and do not achieve proportional dominance in the populations of women we have examined to date. Elucidating the ecological mechanisms responsible for this differential “success” of *Gardnerella* spp. will require further focused study.

### Co-occurrence of *Gardnerella* spp

Given the frequency with which women are colonized by multiple species of *Gardnerella*, we were interested to determine if there are any consistent patterns of co-occurrence among species. Closely related species occupying similar environmental niches might be expected to co-occur more frequently, and depending on resource levels, they might also compete with each other. Raw correlations of read counts are not recommended for assessing co-occurrence as they are biased in the context of the compositional nature of amplicon sequence analysis [35–37]. Methods based on presence/absence, such as Jaccard’s index can be informative in this context and perform better than raw correlations [22]. To determine whether taxa co-occur more or less often than expected by chance, a reasonable null model for Jaccard’s index is required. Traditional null models of co-occurrence have used randomizations and simulations, but have been shown to be biased under many circumstances [38]. Therefore, we used a probabilistic null model of co-occurrence [23], which performs well for microbial sequencing data [22]. In addition to co-occurrence analysis using presence/absence, we also investigated proportionality of species [36] on the center-log ratio transformed read counts using the ‘propr’ package [39]. The results were very similar with *G. vaginalis* and *G. swidsinksii* showing clustering by proportionality, as well as *G. piotii* and genome species 3. However, as there is currently no agreed upon hypothesis testing method for proportionality, we used the presence/absence data to determine significant co-occurences.

Significant co-occurrences were observed for several pairs of species, but there were also many cases where species co-occurred only randomly (Figure 4A, Table S3). Among the most frequently detected species (*G. vaginalis*, *G. swidsinskii*, *G. leopoldii*, *G. piotii* and genome sp. 3), the smallest pairwise Jaccard distances (i.e. the most samples in common) were observed for *G. vaginalis* and *G. swidsinskii*, and *G. piotii* and genome sp. 3. (Figure 4B). *G. leopoldii* and *G. swidsinskii* did not occur together more often than expected by chance, which is of note as both were previously labeled as subgroup A based on cpn60 sequences and whole genome sequence comparisons [8]. The differentiating features of these two species detected by MALDI-TOF [13] may be associated with their occupation of distinct niches. This suggests that the new labeling is indeed useful for understanding differences in distributions at this deeper level. None of the species pairs had fewer co-occurrences than expected, suggesting that competitive exclusion may not be important for describing their relative distributions. Conclusions regarding the rarely detected genome species are limited since very few samples were positive and thus chances of observing co-occurrence were correspondingly low.

**Figure 4.**
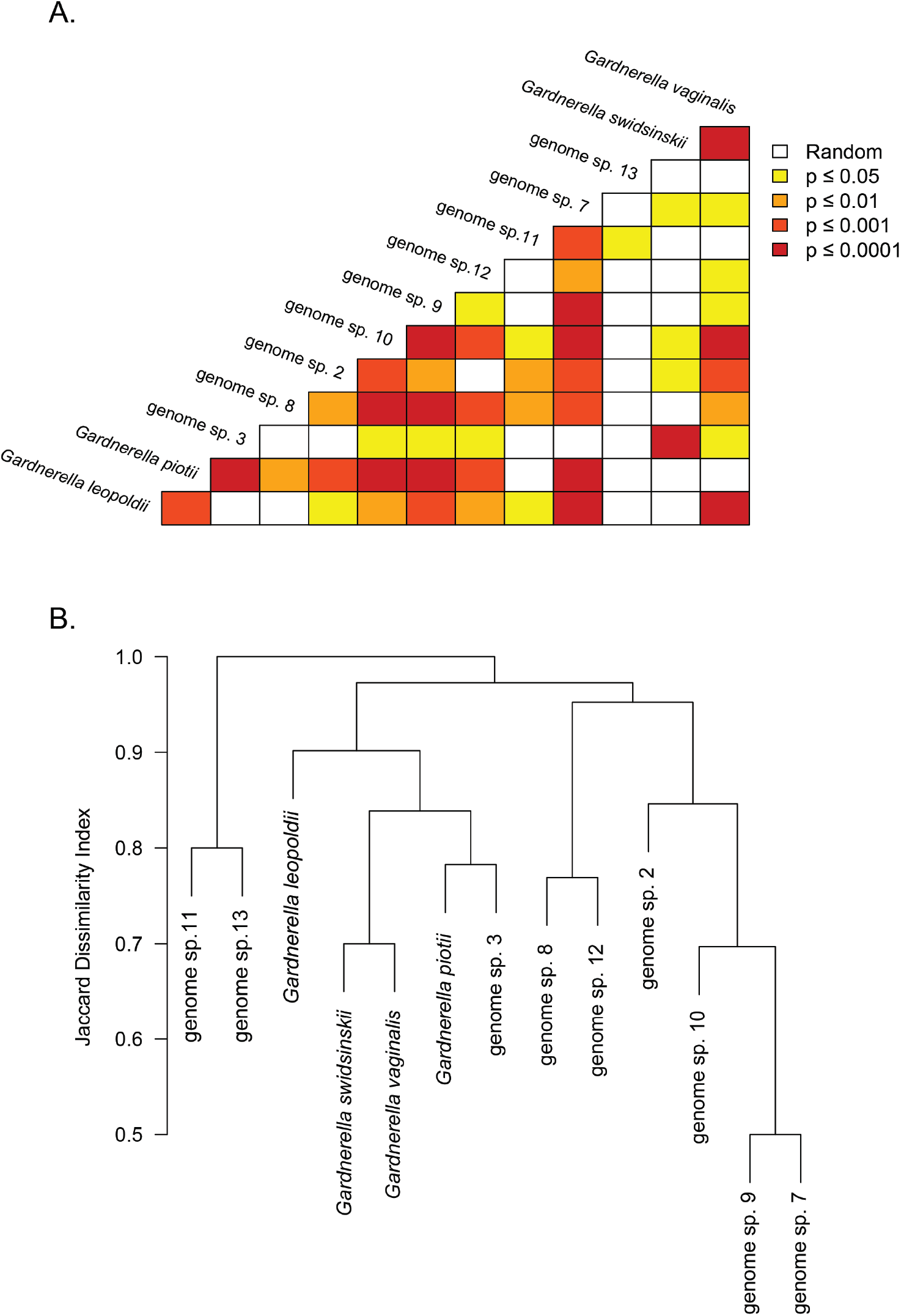
(A) Significance of pairwise co-occurrences of *Gardnerella* spp. in 413 vaginal samples determined by Jaccard index of similarity (*J*) calculation. Size of the *P* value is indicated by colour according to the legend. Species were considered present if the raw number of sequence reads in the sample was ≥10, otherwise the species was marked as absent. *P*-values were Benjamini-Hochberg corrected to a false-discovery rate = 0.05 (B) Hierarchical clustering of species based on Jaccard distances (1 − *J*), using complete linkage.

### Association of *Gardnerella* spp. with BV status and vaginal symptoms

To understand how resolution of different *Gardnerella* spp. may inform clinically important outcomes, we compared the relative abundances of the more frequently occurring species (*G. vaginalis*, *G. swidsinskii*, *G. leopoldii*, *G. piotii* and genome sp. 3) among groups based on clinical Nugent scores (Negative, Intermediate, BV), and self-reported symptoms in the two weeks prior to the swab collection (odour, irritation, and discharge). There was a significant association between Nugent category and relative abundance of *G. vaginalis*, *G. swidsinskii*, and *G. piotii*, (Table 1). Genome sp. 3 was marginally associated with Nugent category, but none of the pairwise comparisons was significant after p-value adjustment. The relationship between *Gardnerella* abundance and Nugent score is not surprising, as the presence of *Gardnerella* “morphotypes” on Gram stained slides of vaginal specimens is part of the calculation of the clinical score [3]. The lack of association of *G. leopoldii* with Nugent category is interesting in that this species was previously investigated together with *G. swidsinskii* as cpn60 subgroup A/clade 4 which was found to be associated with Nugent category [14, 34, 40]. These species are very closely related phylogenetically (Figure 1), and were only resolved by Vaneechoutte et al. [13] by MALDI-TOF and whole genome comparison, so the specific factors responsible for their apparently different relationships with the microbiological definition of BV remain to be identified.

**Table 1.**
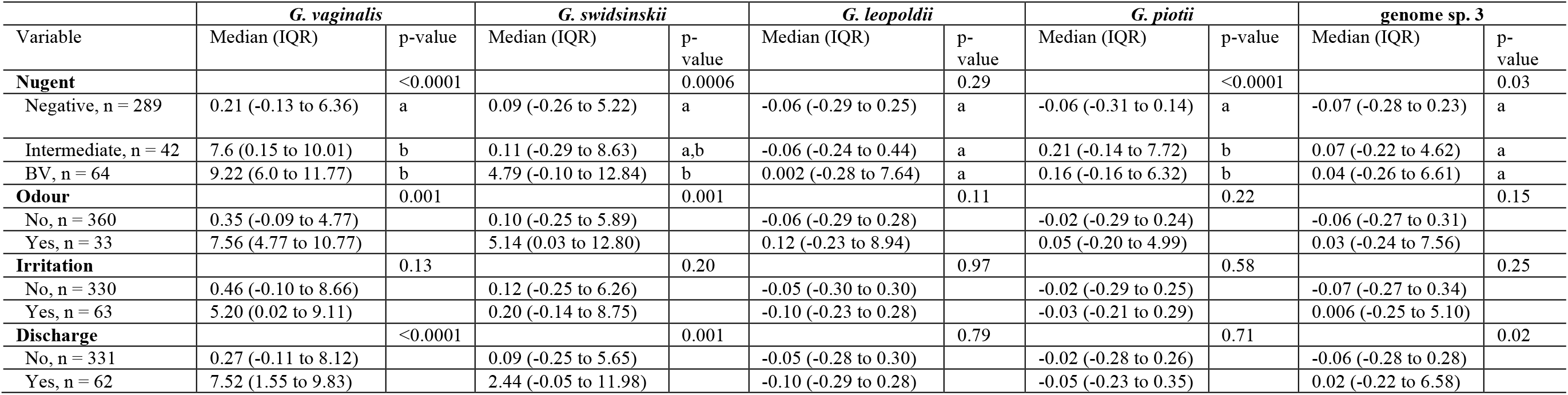
Centre-log-ratio (CLR) transformed relative abundance by clinical variables. P-values are from Kruskal-Wallis tests. Pairwise comparison significance for Nugent category is indicated using letters (a, b) from Dunn tests with Benjamimin-Hochberg p-value adjustment. CLR transform was relative to the geometric mean log2 relative abundance of all taxa (including non-*Gardnerella*) therefore negative values indicate relative abundance less than the mean log2 relative abundance of all taxa.

Phenotypic diversity has long been considered a possible explanation for the detection of *Gardnerella* in women without vaginal symptoms, however, attempts to identify associations between particular biotypes and clinical status have yielded inconsistent and often contradictory results [41–45]. The major limitation of investigations relying on phenotypic characterization of isolates is that they focus only on the most readily culturable isolates from individual specimens (often only one isolate per specimen), which is inadequate since women are usually colonized by multiple species of *Gardnerella*. We observed strong relationships between abnormal odour and discharge with higher relative abundance of *G. vaginalis* and *G. swidsinskii*, but not with the other three species, although there is a marginal relationship between discharge and genome sp. 3 (p = 0.02) (Table 1). *G. vaginalis* and *G. swidsinskii* co-occurred more often than expected by chance, and also showed proportionality suggesting that they are correlated in abundance. Therefore, we cannot be sure if it is just one species or both that is associated with vaginal symptoms. Sialidase activity defines *G. piotii* [13] and is also observed for genome sp. 3 isolates [8], however, these species were not associated with discharge nor were they the most strongly associated with a BV diagnosis by Nugent score. This is likely due to the polymicrobial nature of BV, and the fact that many other BV-associated bacteria produce hydrolytic enzymes that may contribute to symptoms [46–48]. The lack of association of sialidase positive *Gardnerella* spp. with symptoms is also consistent with the suggestion that some types of *Gardnerella* may be important for “stage-setting”; establishing an anaerobic environment and initial adhesion to the vaginal epithelium that lead to abundant growth of other BV associated organisms, and the development of multi-species biofilms (reviewed in [49]). In these primary colonizers, hydrolytic enzymes and cholesterol-dependent cytolysin (vaginolysin) may be more important in preparing the microbiome for secondary expansion of populations of BV associated bacteria rather than acting as specific virulence factors affecting the host.

## Conclusions

Considering *Gardnerella* as a monolithic taxon in vaginal microbiome studies (due to the ubiquitous application of 16S rRNA gene sequencing in microbiome profiling) has limited progress in understanding the link between vaginal microbiota and clinical outcomes, and the development of improved diagnostics for women’s health. Our results provide a clear demonstration of the utility of cpn60 barcode sequencing for rapid, high-throughput determination of *Gardnerella* spp. abundance and distribution in the vaginal microbiome with minimal sequencing effort. This approach will be critical in further investigation of the intriguing association of *G. piotii* (subgroup B/clade 2) with “intermediate” microbiota, which has been observed independently using cpn60 barcode sequencing and clade-specific PCR [14, 33]. It remains to be determined if *Gardnerella* species that do not regularly achieve numerical dominance in the microbiome contribute to establishment and maintenance of dysbiosis. Robust and simple classification of *Gardnerella* isolates based on cpn60 barcode sequences will facilitate further characterization of these organisms within the new taxonomic framework, and lead to identification of phenotypic features of the species that determine their ecological roles in the vaginal microbiome. In the clinical context, assessing longitudinal shifts in *Gardnerella* spp. abundance will also be important to evaluate natural or post treatment changes. In future cohort studies, application of cpn60 barcode sequencing will provide new insight as to whether *Gardnerella* spp. diversity and differential distribution are an explanation for issues such as treatment failure and recurrent vaginal dysbiosis [34, 50], or for the failure of antimicrobial treatment in the prevention of preterm birth despite a strong association between vaginal dysbiosis and preterm delivery [51].

## Declarations

### Ethics approval and consent to participate

Studies from which data was accessed for this sub-study were approved by the University of British Columbia Children’s & Women’s Research Ethics Board (Certificate numbers H10-02535, H11-00119, and H11-01912).

### Availability of data and materials

The datasets supporting the results of this article is available in the NCBI repository (BioProject Accessions: PRJNA362575, PRJNA278895, PRJNA528096). R code for the co-occurrence analysis and data table are provided as supplemental information.

### Authors’ contributions

JEH and AYKA conceived the study, conducted the analysis and wrote the paper. The VOGUE Research Group provided access to data for analysis and edited the paper. All authors read and approved the final manuscript.

## Acknowledgements

The VOGUE Research Group is Deborah Money, Alan Bocking, Sean Hemmingsen, Janet Hill, Gregor Reid, Tim Dumonceaux, Gregory Gloor, Matthew Links, Kieran O’Doherty, Patrick Tang, Julianne van Schalkwyk and Mark Yudin.

## Supplemental Information

**Table S1.** Pairwise DNA sequence identities among *Gardnerella* spp. based on a CLUSTALw alignment of the 552 bp cpn60 barcode sequence.

**Table S2.** Sequence read frequencies for 13 *Gardnerella* species in 413 vaginal microbiome samples.

**Table S3.** Numbers of occurrences and co-occurrences, expected co-occurrences and *P* values for pairwise comparisons of 13 *Gardnerella* species in 413 vaginal samples.

**Supplemental File 1.** R code for co-occurrence analysis.

